# A novel flatworm-specific gene family implicated in reproduction in *Macrostomum lignano*

**DOI:** 10.1101/167346

**Authors:** Magda Grudniewska, Stijn Mouton, Margriet Grelling, Anouk H. G. Wolters, Jeroen Kuipers, Ben N. G. Giepmans, Eugene Berezikov

**Author notes:** These authors contributed equally to this work (Magda Grudniewska, Stijn Mouton).

## Abstract

Free-living flatworms, such as the planarian *Schmidtea mediterranea*, are extensively used as model organisms to study stem cells and regeneration. The majority of studies in planarians so far focused on broadly conserved genes. However, investigating what makes these animals different might be equally informative for understanding its biology. Here, we present a re-analysis of neoblast and germline transcriptional signatures in the flatworm *M. lignano* and combine it with the whole-animal electron microscopy atlas (nanotomy) as a reference platform for ultrastructural studies in *M. lignano*. We show that germline-enriched genes have a high fraction of flatworm-specific genes and identify *Mlig-sperm1* gene as a member of a novel gene family conserved only in free-living flatworms and essential for producing healthy spermatozoa. This work demonstrates that investigation of flatworm-specific genes is crucial for understanding flatworm biology and establishes a basis for future research in this direction in *M. lignano*.

## Introduction

Animal models inspired researchers for hundreds of years. In biomedicine, a wide range of organisms is employed to study e.g. development, ageing, and mechanistic underpinnings of diseases, with the aim of translating these findings to humans. Sometimes, however, it is the unique feature of the experimental model that brings the breakthrough. A great example is the discovery of the angiotensin-converting enzyme (ACE) inhibitor. This chemical molecule derived from the Brazilian viper venom, was successfully used in the development of a drug to treat hypertension and acute myocardial infarction (Komajda and Wimart, 2000). Another compound, squalamine, isolated from dogfish sharks, exhibits strong anti-fungal and anti-bacterial activity. It was shown to be very efficient in fighting a broad spectrum of human pathogens, strengthening its therapeutic potential (Zasloff et al., 2011). Finally, a more recent study deciphering the remarkable resistance of tardigrades to X-ray radiation, led to the discovery of a novel DNA protector, the Dsup protein. When expressed in human cells, this protein seems to have the ability to protect human DNA as well (Hashimoto et al., 2016).

Given the above examples of successful, therapeutic use of the unique characteristics of some animals, and the large number of organisms demonstrating extreme features, such as a remarkable resistance to environmental factors, astonishing regeneration abilities, or an extremely long lifespan, it is essential to explore the potential of nontraditional models for biomedicine (Austad, 2009, 2010; De Magalhães, 2015; Valenzano et al., 2017).

One of such organisms is the free-living flatworm *Macrostomum lignano*, which has been developed into an experimental platform to study various biological phenomena (Arbore et al., 2015; Rivera-Ingraham et al., 2016; Vellnow et al., 2017; Wasik et al., 2015). Several experimental protocols have been established for this animal, including transgenesis methods (Wudarski et al., 2017). Furthermore, we recently characterized gene expression in the proliferating cells of *M. lignano* and established transcriptional signatures of proliferating somatic neoblasts, including the stem cells, and germline cells and demonstrated the role of several genes conserved between *M. lignano* and human in stem cell and germline biology (Grudniewska et al., 2016). In the present follow-up study, we reanalyzed the published dataset using an improved transcriptome assembly and focused on genes that are not conserved in human. We demonstrate that knockdown of one of the tested non-conserved genes can lead to aberrations in gonad and sperm structure. We advocate that functional studies of non-conserved genes in *M. lignano* will be crucial for understanding the biology of this model organism and may lead to discoveries translatable to human health, such as improved wound healing or novel anthelminthic drugs.

## Results and Discussion

### Characterization of conservation levels in different gene groups

All previous RNA-seq based gene expression studies in M. lignano relied on de novo transcriptome assemblies (Arbore et al., 2015; Grudniewska et al., 2016; Wasik et al., 2015). We have recently generated a significantly improved genome assembly for the *M. lignano* DV1 line (Wudarski et al., 2017), which in turn allowed us to generate a genome-guided transcriptome assembly Mlig_RNA_3_7_DV1_v3. The new transcriptome assembly resolved many partial transcript issues inherent for de novo transcriptomes (Wudarski et al., 2017). We re-analyzed the previously established proliferating neoblast and germline transcriptional signatures (Grudniewska et al., 2016) using the new transcriptome assembly (**Supplementary Table 1, Supplementary Figure 1**). While the conclusions of the previous work do not change with the new analysis, the number of transcript clusters classified as ‘stringent neoblast’ increased from 357 to 489 (**Supplementary Figure 1K**), while the number of ‘irradiation’ and ‘germline’ transcript clusters decreased from 7,277 to 5,901 and from 2,739 to 2,604 respectively (**Supplementary Figure 1C, G**). These changes are explained by the less-fragmented transcriptome assembly, which also allows more accurate estimation of gene conservation depth. We assigned the conservation level to each M. lignano gene as ‘conserved’ if there is a significant homology with human genes, as ‘flatworm-specific’ if homologs are identified only in *Schmidtea mediterranea* and/or *Schistosoma mansoni*, or as ‘*M. lignano*-specific’ if no homologs are detected (**Table 1**). The distribution analysis of the conservation levels between different gene categories revealed striking differences between neoblast and germline genes. While overall 47.3% of M. lignano genes are conserved in human, 8.2% are flatworm-specific and 44.5% are *Macrostomum*-specific (**Table 1**), for the neoblast genes the fraction of human-conserved genes is substantially higher at 85%, while flatworm-specific and non-conserved gene fractions are only 2.8% and 12.2% respectively (**Table 1**). At the same time, the fraction of germline genes conserved in human is 37.6%, which is significantly less than overall, while the fraction of *Macrostomum*-specific genes rises to 54.2% (**Table 1**). Since in this analysis we used all transcripts from the Mlig_RNA_3_7_DV1_v3 transcriptome assembly, including transcripts without predicted open reading frame (ORF), it is possible that the fraction of *Macrostomum*-specific transcripts is inflated. We repeated the conservation distribution analysis using only transcripts with ORFs and clustering sequences at 95% amino-acid identity level to exclude biases due to possible expansions of gene families. However, the picture did not change significantly: overall 55.3% of genes are conserved in human, 9.7% are flatworm-specific and 35% are *Macrostomum*-specific, while the numbers are 86%,3 % and 11% for the neoblast genes and 42.5%, 9% and 48.5% for the germline genes respectively (**Table 1**). The observed differences in conservation levels between neoblasts and germline are in line with published literature on deep conservation of the neoblast regulation program (Önal et al., 2012) and significant variation in evolution of reproductive systems (Schwander et al., 2014).

**Table 1.** Conservation of different *M. lignano* gene groups in human and flatworms.

|  | Total number | Homologs in human | Homologs in *S. mediterranea* | Homologs in *S. mansoni* | Flatworm-specific* | *M. lignano*-specific |
| --- | --- | --- | --- | --- | --- | --- |
| **Transcript clusters** |  |  |  |  |  |  |
| All | 50,673 | 23,969<br>(47.30%) | 25,716<br>(50.75%) | 21,581<br>(42.59%) | 4,159<br>(8.21%) | 22,545<br>(44.49%) |
| Neoblasts | 1,062 | 902<br>(84.93%) | 871<br>(82.02%) | 838<br>(78.91%) | 30<br>(2.82%) | 130<br>(12.24%) |
| Neoblasts, stringent | 489 | 406<br>(83.03%) | 379<br>(77.51%) | 370<br>(75.66%) | 14<br>(2.86%) | 69<br>(14.11%) |
| Germline | 2,604 | 979<br>(37.60%) | 1067<br>(40.98%) | 840<br>(32.26%) | 213<br>(8.18%) | 1,412<br>(54.22%) |
| **Protein clusters** |  |  |  |  |  |  |
| All | 30,017 | 16,605<br>(55.32%) | 17,758<br>(59.16%) | 14,819<br>(49.37%) | 2,913<br>(9.70%) | 10,499<br>(34.98%) |
| Neoblasts | 840 | 723<br>(86.07%) | 693<br>(82.50%) | 676<br>(80.48%) | 25<br>(2.98%) | 92<br>(10.95%) |
| Neoblasts, stringent | 404 | 336<br>(83.17%) | 310<br>(76.73%) | 307<br>(75.99%) | 10<br>(2.48%) | 58<br>(14.36%) |
| Germline | 1,917 | 815<br>(42.51%) | 882<br>(46.01%) | 707<br>(36.88%) | 173<br>(9.02%) | 929<br>(48.46%) |

\* present in *S. mediterranea* or *S. mansoni* but not in human.

### Knockdown of a flatworm-specific gene *Mlig-sperm1* causes abnormal morphology of testes and spermatozoa and decreased fertility

To assess the possible roles of non-conserved and flatworm-specific genes in *Macrostomum* biology, we randomly chose six candidate genes enriched in proliferating cells for a pilot functional screen (**Supplementary Table 2**). We carried out knockdown experiments, including two conditions: homeostasis and regeneration. From all the tested candidates, one of the germline genes, *Mlig020950*, demonstrated a strong phenotype. We named the *Mlig020950* gene as *Mlig-sperm1* due to severe defects in sperm morphology in *Mlig020950(RNAi)* animals, as described below.

Gene knockdown of *Mlig-sperm1* led to an aberrant morphology of testes, which were dramatically enlarged in all individuals (**Figure 1A-D**). Detailed analysis revealed that the oversized testes accumulated large amounts of sperm cells (**Figure 1E-F**). The spermatozoa of *Mlig-sperm1(RNAi)* individuals demonstrated aberrant morphology (teratozoospermia), indicated by a very rigid shaft, and often forming contortion at the notch site (**Figure 1F**). In contrast to sperm of control animals, which demonstrate undulating movements of the shaft and distal process (**Supplementary Video 1**), knockdown worms’ spermatozoa showed atypical motility (asthenozoospermia): cells were not swimming actively and performed twitching movements (**Supplementary Video 2**).

**Figure 1.**
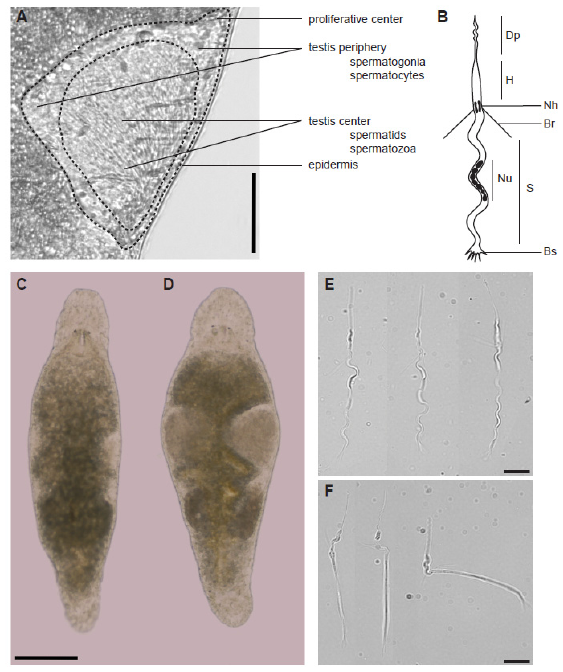
The structure of testis and sperm cell in healthy control and *Mlig-sperm1* knockdown *M. lignano* animals. (A) The structure of the testis. The testis periphery contains a number of spermatogonia and primary and secondary spermatocytes. The anterior tip of the testes is described as ‘proliferative center’. The inner region of testis is filled by maturing spermatids and mature spermatozoa. The latter are discharged into the vas deferens at the posterior end of the testes. Sperm is then stored in the seminal vesicles within the tail plate. (B) Schematic structure of an adult sperm cell. Dp, distal process (feeler); H, head; Nh, notch; Br, bristle; Nu, nucleus; S, shaft; Bs, brush. For a detailed description of sperm cells in *M. lignano* see Willems et al., 2009. (C, D) Comparison of overall morphology between control (C) and *Mlig-sperm1(RNAi)* (D) individuals. (E, F) Comparison of sperm morphology between control (E) and knockdown (F) worms using DIC microscopy. Scale bars are 100 µm (A, C, D) and 10 µm (E, F).

Scanning electron microscopy (EM) confirmed the morphological aberrancy of *Mlig-sperm1(RNAi)* spermatozoa (**Figure 2**). In comparison to control cells (**Figure 2A-C**), the shaft of the *Mlig-sperm1* knockdown spermatozoa, demonstrates rigidity and lack of curvature (**Figure 2D**), and its brush is flattened and inflexible (**Figure 2E**). Furthermore, the contortion at the notch site is clearly visible (**Figure 2F**). In addition, scanning transmission EM demonstrated that also the nuclei of late spermatids and spermatozoa of *Mlig-sperm1(RNAi)* worms have an aberrant morphology (**Figure 3**). In controls, the nucleus elongates and chromatin condenses into a number of discrete bodies connected by small bridges during the late phase of spermiogenesis (**Figure 3C-E**) as previously described (Willems et al., 2009). In *Mlig-sperm1(RNAi)* worms, distinct nuclear bodies can be rarely observed and the chromatin has a fragmented appearance (**Figure 3F-H**).

**Figure 2.**
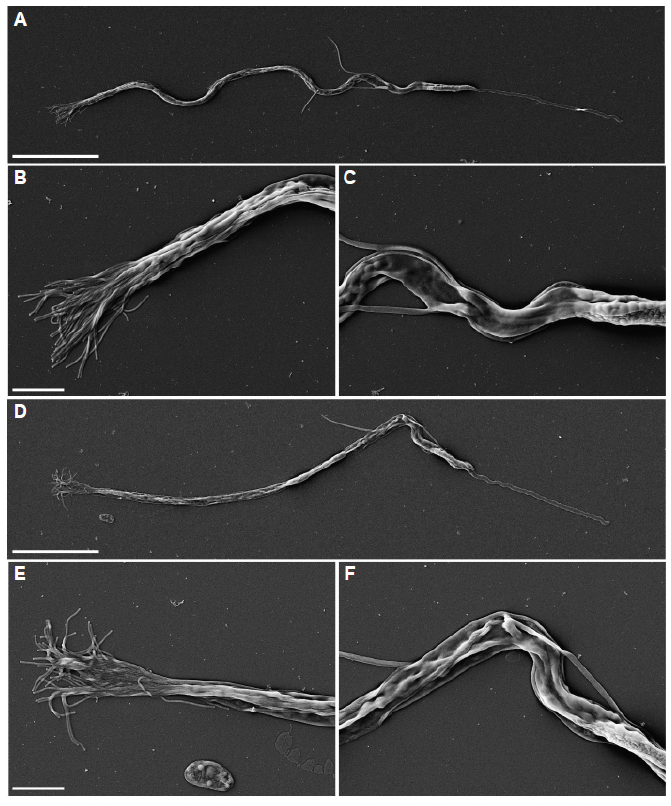
Scanning electron microscopy of spermatozoa. (A-C) Spermatozoon of a negative control *GFP(RNAi)* worm. (A) Overview of the complete cell. Note the curved view of the shaft. (B) Detail of the brush, consisting out of separate extensions. (C) Detail of the notch region of the cell. (D-F) Spermatozoon of a *Mlig-sperm1(RNAi)* worm. (D) Overview of the complete cell. Note the rigidity of the shaft and the contortion at the notch site. (E) Detail of the brush. Compared to the negative control the brush looks more flattened with the base of the extensions being more packed together. (F) Detail of the notch region clearly showing the contortion. Scale bar are 10 µm (A, D) and 2 µm (B, C, E, F).

As sperm defects are often reported as a cause of decreased male fertility or even complete sterility (Wosnitzer et al., 2014; Yatsenko et al., 2010), we compared fertility of control and *Mlig-sperm1(RNAi)* animals. The knockdown of *Mlig-sperm1* gene resulted in a significantly lower (p < 0.05, *t-test*) number of progeny in each of the three studied weeks of the RNAi treatment (**Supplementary Figure 2**) Interestingly, the number of hatchlings was further decreased during the third week of the experiment compared to the first two weeks (**Supplementary Figure 2**). We speculate that the effects of gene knockdown may strengthen with time, potentially resulting in sterility after a sufficient period of dsRNA treatment.

### *Mlig-sperm1* is expressed exclusively in testes

*Mlig-sperm1* is categorized as enriched in proliferating germline cells according to the RNA-seq data (**Supplementary Table 1**), and to confirm the gene expression pattern we performed whole mount *in situ* hybridization (WISH), including 1-day old hatchlings, 4-5 days old juveniles and adult individuals. Based on light microscopic analysis, no signal was detected in hatchlings (**Supplementary Figure 3A**), suggesting that the gene is not expressed in the gonad anlage of primordial germ cells (Pfister et al., 2008). In the juveniles, the signal was observed on the level of developing testes in a cluster of several cells (**Supplementary Figure 3B**). Finally, a strong expression of *Mlig-sperm1* was detected in adult animals, in the cells located in the testes periphery, known as spermatogonia and/or spermatocytes (**Supplementary Figure 3C** and **Figure 1A**). Specificity of ISH labelling was confirmed by a *Mlig-sperm1* sense probe, for which no signal was detected (**Supplementary Figure 3D**).

Fluorescent *in situ* hybridization (FISH) and confocal microscopy analysis of adult worms confirmed the pattern observed with WISH (**Supplementary Figure 3E, F**). In addition, FISH revealed expression of *Mlig-sperm1* in the tip of the testes, sometimes referred to as ‘the proliferative center’ of the testes (**Supplementary Figure 3E, arrowhead**, and **Figure 1A**). These results correspond to the initial classification of *Mlig-sperm1* as a gene enriched in proliferating germline cells. We did, however, also detect signal of the gene in spermatozoa discharging into the vas deferens (**Supplementary Figure 3F, arrowhead**), which suggest that *Mlig-sperm1* is continuously expressed in the later stages of spermatogenesis. In the light of these findings, we propose that the *Mlig-sperm1* gene is expressed in the proliferating and differentiated compartment of male gonads and plays a role in forming healthy sperm in *M. lignano*. Its exact function is, however, yet to be determined.

### *Mlig-sperm1* is a member of a large free-living flatworm-specific gene family

The *Mlig-sperm1* gene has two nearly identical loci in the Mlig_3_7_DV1 genome assembly, *Mlig020950.g1* and *Mlig020950*.g2. We next studied the relation of this gene to other *Macrostomum* genes and homologs in other flatworms. The BLAST search revealed that *Mlig-sperm1* has a significant similarity with 35 genes in *M. lignano* (blastp e-value cutoff 1e-15), which can be grouped into 14 protein clusters based on 95% amino-acid identity cutoff (**Supplementary Table 3**). Notably, most of the homolog genes are enriched in the proliferating germline cells according to the previous analysis (Grudniewska et al., 2016). Furthermore, the gene is conserved in *S. mediterranea*, with one and 8 transcripts identified in asexual (dd_Smed_v6) and sexual (dd_Smes_v1) transcriptome assemblies respectively (**Supplementary Table 3**), as well as in all 5 other planarian species available in PlanMine (Brandl et al., 2015). We did not find significant matches to the Mlig-SPERM1 protein in the transcriptomes of parasitic flatworms or outside flatworm species. Alignment of the representative proteins revealed a conserved domain common to all genes (**Supplementary Figure 4**). However, a search against the Pfam database did not reveal homology to any known protein family. The tertiary structure of the Mlig-SPERM1 protein, as predicted by RaptorX (Peng and Xu, 2011), consists of three domains with 48% of the predicted positions being disorganized. The region conserved between all the genes (**Supplementary Figure 4**) corresponds to the first domain, which has the highest organization factor and consists of one beta sheet and several alpha helixes (**Supplementary Figure 5**). Given the conservation level between *Mlig-sperm1* and other genes, and the classification of many of the genes as enriched in proliferating germline cells, we suggest that *Mlig-sperm1* is a member of a novel protein family specific for free-living flatworms, with important roles in flatworm reproduction.

### *M. lignano* nanotomy resource

The transmission electron microscopy analysis of control and *Mlig-sperm1(RNAi)* animals presented in **Figure 3** was performed using the anatomy at the nanoscale (nanotomy), approach, which allows scanning of large specimen areas (Ravelli et al., 2013) and provides a ‘Google-Earth’ style of data presentation and navigation at different levels of resolution. In this study images of longitudinal sections of whole animals under *Mlig-sperm1(RNAi)* and *GFP(RNAi)* conditions were generated (**Supplementary Figure 6**). Furthermore, for a wild-type worm we generated 35 cross-sections covering a complete animal (**Supplementary Figure 6**). The generated resource is available at http://www.nanotomy.org/OA/Macrostomum. While the detailed annotation of these nanotomy images is beyond the scope of this work, we believe that the generated resource will serve as a valuable reference on *M. lignano* morphology at ultrastructural level and complements genomic resources available for this developing model organism.

**Figure 3.**
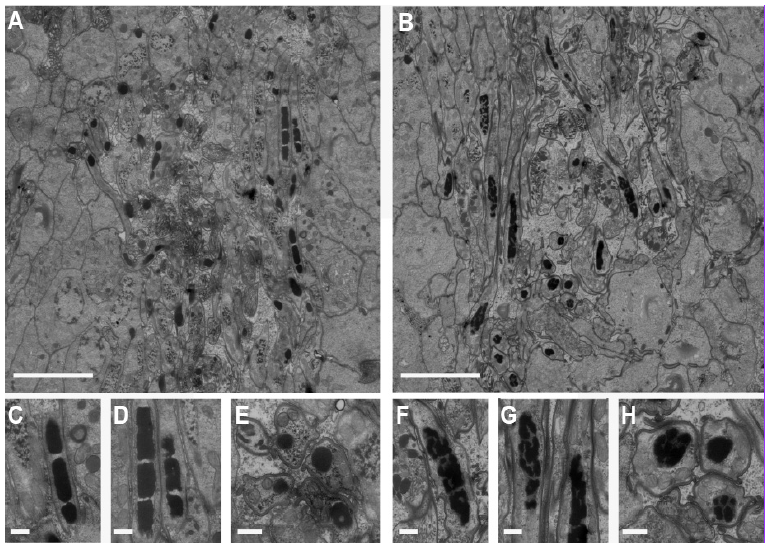
Ultrastructure of spermatids and spermatozoa. (A, B) Overview of early and late spermatids and spermatozoa in the testes of a negative control *GFP(RNAi)* worm (A) and a *Mlig-sperm1(RNAi)* worm (B). In both images, several longitudinal and cross sections of nuclei of the spermatozoa can be observed as black structures. (C, D) Detail of longitudinal sections of spermatozoa nuclei of a *GFP(RNAi)* worm. The chromatin of the nucleus is condensed into discrete bodies. (E) Detail of a cross section of spermatozoa nuclei of a *GFP(RNAi)* worm. (F-G) Detail of longitudinal sections of spermatozoa nuclei of a *Mlig-sperm1(RNAi)* worm. Compared to the negative control, the chromatin of the nuclei looks fragmented and condensation into discrete bodies is less visible. (H) Detail of a cross section of spermatozoa nuclei of a *Mlig-sperm1(RNAi)* worm. Compared to the negative control, the chromatin of the nuclei is more fragmented. Scale bars are 5 µm (A, B) and 500 nm (C-H).

## Conclusions

In the present study, we reanalyzed the transcriptomic signatures of proliferating cells in *M. lignano* using an improved transcriptome assembly and investigated the extent of conservation of *M. lignano* genes with neoblast- and germline-enriched expression. Neoblast-enriched genes were found to be deeply conserved, while germline-enriched genes have many non-conserved or flatworm-specific genes. Moreover, we demonstrated that a novel gene, which we named *Mlig-sperm1*, is expressed in all cell types of the testes and is essential during spermatogenesis. This example illustrates that our approach of selecting non-conserved or flatworm-specific genes can yield important insight into aspects of flatworm biology, such as germline development. Future studies of genes conserved within flatworms, and more specifically in parasitic flatworms, could help to develop treatments for infections caused by parasites, such as *Schistosoma*. Current research in that area indeed focuses its efforts on dissecting the mechanism behind the maintenance and activity of the germline, as the egg-induced inflammation is the main cause of *Schistosoma*-associated pathologies (Collins & Newmark, 2013; Iyer, Collins, & Newmark, 2016).

Focusing on flatworm-specific genes enriched in the neoblasts could contribute to our understanding of e.g. their astonishing regeneration capacity and would create an opportunity to improve such competencies in humans. This is in line with the recent report on improved radiotolerance of human cultured cells by a tardigrade-unique protein (Hashimoto et al., 2016). We therefore advocate that investigating *M. lignano* genes not conserved in humans is an approach with truly great potential.

## Materials and Methods

### Experimental organism and culture conditions

*Macrostomum lignano* (Macrostomida, Rhabditophora) is a free-living marine flatworm and an obligatory, non-self-fertilizing hermaphrodite reproducing exclusively in a sexual manner. (Rieger et al., 1988). The combination of a short generation time of about 3 weeks and high fertility rates allows a rapid expansion of cultures (Ladurner et al., 2008).

The animal is small, about 1 mm long and consists of approximately 25,000 cells (Ladurner et al., 2000). Worms are transparent and major tissues and organs can be easily distinguished (i.e. eyes, brain, gonads, reproductive organs, gut). Worms are kept in Petri dishes with nutrient-enriched artificial seawater (f/2) (Anderson et al., 2005), at 20°C and a 14/10 hours light/dark cycle and are fed *ad libitum* with the diatom *Nitzschia curvilineata* (Rieger et al., 1988).

In the present study, the recently collected and introduced in the Berezikov lab, NL10 strain was used. In contrast to DV1 (Zadesenets et al., 2016), NL10 does not demonstrate chromosomal polymorphism (Wudarski et al., 2017).

### Conservation and alignments

Amino-acid sequences from the Mlig_RNA_3_7_DV1_v3 transcriptome assembly (Wudarski et al., 2017) were queried against human, *S. mediterranea* (dd_Smed_v6 and dd_Smes_v1) and *S. mansoni* (ASM23792v2) genes using BLAST (Altschul et al., 1997) and hits with E-value below 0.01 were considered as homologs for the purpose of conservation analysis. Sequence alignment and visualization were performed with CLC Main Workbench (QIAGEN Aarhus A/S).

### Whole mount *in situ* hybridization

cDNA synthesis was performed using the SuperScript III First-Strand Synthesis System (Life Technologies) according to the manufacturer’s protocol with 2–3 µg of total RNA as a template for each reaction. Provided oligo(dT) and hexamer random primers were used.

DNA fragments selected as templates for *in situ* hybridization probes, were amplified from cDNA by standard PCR with GoTaq Flexi DNA Polymerase (Promega), followed by cloning using the pGEM-T vector system (Promega) and sequenced by GATC Biotech. All primers used are listed in Supplementary table 3. DNA templates for producing DIG – labeled riboprobes were amplified from sequenced plasmids using High Fidelity Pfu polymerase (Thermo Scientific). Forward (5’-CGGCCGCCATGGCCGCGGGA-3’) and reversed (5’TGCAGGCGGCCGCACTAGTG-3’) primers binding the pGEM-T vector backbone near the insertion site were designed. Moreover, versions of the same primers with a T7 promoter sequence (5’-GGATCCTAATACGACTCACTATAGG-3’) appended upstream were obtained. The T7 promoter sequence serves as a start site in subsequent in vitro transcriptions. A pair of primers, depending on the orientation of the insert in the vector: forward with T7 promoter and reverse without or vice versa, was used to amplify every ISH probe template.

Digoxigenin (DIG) labeled RNA probes (800 to 1200 bp in length) were generated using the DIG RNA labeling Mix (Roche, Switzerland) and T7 RNA polymerase (Promega, Fitchburg, WI) according to the manufacturer’s protocol for in vitro transcription. The concentration of every probe was measured with the Qubit RNA BR assay (Invitrogen), probes were diluted in Hybridization Mix (Pfister et al., 2007) to 20 ng/µl, stored at −80°C and used within 4 months. The final concentration of the probe and optimal temperature used for hybridization varied for different probes and were optimized for each probe.

Whole mount in situ hybridization (ISH) was performed following published protocol (Pfister et al., 2007). Pictures were made using a standard light microscope with DIC optics and an AxioCam HRC (Zeiss, Germany) digital camera and the EVOS XL Core Imaging System (ThermoFisher).

### Fluorescent *in situ* hybridization and immunofluorescence

Fluorescent in situ hybridization (FISH) was performed following the published FastBlue protocol developed for planarians (Currie et al., 2016), except the 5% NAC treatment and bleaching steps were omitted. Slides were mounted using 80% glycerol solution, and the labeling was visualized with a Leica TCS SP8 confocal microscope at the UMCG Imaging and Microscopy Center.

### RNA interference

In order to generate dsRNA fragments, the same plasmids were used as for making ISH probes. Templates for the synthesis of both sense and antisense RNA strands were amplified from the plasmids containing the insert of interest. The same primers were used as for ISH riboprobe template amplification, and for each fragment, two PCRs were performed – with both pairs of primers (forward with T7 promoter and reversed without and vice versa). High Fidelity Pfu polymerase (Thermo Scientific) in 150 µl of total volume reaction was used. PCR products were then run on 1% agarose gel, PCR product bands were cut out and purified using the QIAquick Gel Extraction Kit (Qiagen, Netherlands). Each template was then used to synthesize the corresponding single strand RNA with the TranscriptAid T7 High Yield Transcription Kit (Thermo Scientific) according to manufacturer’s protocol. The single reaction volume was 50 µl, and tubes were incubated in 37°C for 5 hours. Afterwards 100 µl of nuclease-free water was added to each tube, sense and antisense RNA strands were mixed to a final volume of 300 µl and annealed by incubating them at 70°C for 10 min, followed by gradual cooling down to room temperature, taking approximately 90 min. Every sample was then treated with 1U of RNase A (Life Technologies) and 5U of DNase I (Thermo Scientific) for 45 min at 37°C. Samples were alcohol precipitated overnight at −80°C. dsRNA was pelleted by centrifugation at 12,000g for 15 min at 4°C, washed with 75% ethanol, and air-dried for 5 min. dsRNA was resuspended in nuclease-free water and the concentration was measured using Nanodrop ND1000. Freshly autoclaved and filtered f/2 medium was used to adjust the concentration to 10 ng/µl. Samples were aliquoted in 1.5 ml Eppendorf tubes and stored at −80°C.

Specific knockdown of candidate genes by RNA interference with double-stranded RNA delivered by soaking was performed as previously described (De Mulder et al., 2009). RNAi soaking experiments were performed in 24-well plates in which algae were grown. Individual wells contained 300 ml of dsRNA solution (10 ng/ml in f/2 medium) in which 15 individuals were maintained. RNAi was performed for three weeks during which dsRNA solution was refreshed daily. Worms were weekly transferred to fresh 24-well plates to ensure sufficient amount of food. For each gene of interest, the effect on homeostasis and regeneration was studied. As a negative control, GFP dsRNA was used. In experiments addressing regeneration, the tail of worms was amputated after 1 week of RNAi. Photos of randomly selected worms were made 1 week after cutting for studying the effect of RNAi on regeneration, and between 2 and 3 weeks of treatment to study to effect on homeostasis.

### Fertility experiment

Worms were treated for three weeks with *Mlig020950* dsRNA or kept in f/2 as a negative control. After that, worms were randomly selected and divided into four groups of five worms. These were cultured in freshly prepared 12-well plates for two weeks while RNAi treatment was ongoing. As a measure of fertility, the number of hatchlings from eggs laid by both experimental and control worms was counted twice a week.

### Electron Microscopy (EM)

*Scanning EM to define surface structure using secondary electron detection (**Figure 2**)* - To isolate spermatozoa, *Mlig020950(RNAi)* and *GFP(RNAi)* worms were relaxed in 7.14% MgCl.6H2O and cut through the testes on a glass slide, using a surgical blade. The cells within the testes were then squeezed out and pipetted onto a poly-l-lysine coated coverslip. After fixation in 2% glutaraldehyde plus 2% paraformaldehyde in 0.1M sodium cacodylate, samples were postfixed with 1% Osmium tetroxide for 30 minutes at 4ºC. Slides were rinsed three times with water, and dehydrated through increasing concentrations of ethanol. Samples were incubated for 10 min in a 1:1 mixture of absolute ethanol and tetramethylsilane on ice, followed by 10 minutes incubation in pure tetramethylsilane on ice. Samples were air dried, glued on aluminium stubs using double sided carbon tape, sputter coated with 10 nm Pd/Au and imaged in a Zeiss Supra55 Scanning Electron Microscope operated at 5 KV using secondary electron detection.

*Scanning transmission EM for ultrastructural analysis of spermatogenesis (**Figure 3**) - Mlig020950(RNAi)* and *GFP(RNAi)* worms were relaxed in 7.14% MgCl.6H2O and fixed in 2% glutaraldehyde plus 2% paraformaldehyde in 0.1M sodium cacodylate buffer for 24 hours at 4°C. After postfixaton in 1% osmium tetroxide/1.5% potassium ferrocyanide for 2 hours at 4°C, worms were dehydrated using ethanol and embedded in EPON epoxy resin. Sections of 60 nm were collected on single slot grids and contrasted using 5% uranyl acetate in water for 20 min, followed by Reynolds lead citrate for 2 min. The longitudinal sections were scanned as described before (Kuipers et al., 2015).

*Scanning EM for transversal nanotomy of Macrostomum (back scatter detector; **Supplementary Figure 6**)*. – Macrostomum lignano was fixed in 2% glutaraldehyde plus 2% paraformaldehyde in 0.1M sodium cacodylate buffer and prepared for EM as described above. Thin sections (~100nm) were collected every 30 µm on silicon wafers as described before (Kuipers et al., 2015b). Data was acquired on a Zeiss Supra 55 scanning EM (SEM) using a back scatter detector (BSD) at 5 kV with 5 nanometer pixel size, 5 µs dwell time using an external scan generator ATLAS 5 (Fibics, Canada) and stitched as described before (Kuipers et al., 2015a; Sokol et al., 2015). After tile stitching the data were exported as an html file and uploaded to the online image database (Fig. 1). Data are available at http://www.nanotomy.org/OA/Macrostomum.

## Author contributions

MGru, SM, EB, Conception and design, Acquisition of data, Analysis and interpretation of data, Drafting the article; MGre, JK, AW, BG, Acquisition and analysis of EM data.

## Acknowledgements

We would like to acknowledge Fleur Broek for the preliminary work within the project. Part of the work has been performed in the UMCG Microscopy and Imaging Center (UMIC), sponsored by ZonMW grant 91111.006. This work was supported by the European Research Council Starting Grant (MacModel, grant no. 310765) to EB.

